# Tetramethylrhodamine self-quenching is a probe of conformational change on the scale of 15 – 25 Å

**DOI:** 10.1101/2024.08.22.609131

**Authors:** Paul Girvan, Liming Ying, Charlotte A Dodson

## Abstract

Tetramethylrhodamine (TMR) is a fluorescent dye whose self-quenching has been used as a probe of multiple biological phenomena. We determine the distance-dependence of self-quenching and place bounds on the timescale of TMR dissociation. Our results validate fluorescence self-quenching as an alternative to FRET and enable future assays to be designed with confidence.

## Introduction

Small-molecule fluorophores covalently attached to biomolecules are often used to create tools and sensors to monitor biological processes *in vitro* and in cells. Rhodamine and its derivative dyes have been popular choices in this context due to their low biological toxicity and cross-reactivity, excitation at visible wavelengths, and high quantum yields. Rhodamines dimerise in aqueous solution, and dimerisation is associated with a characteristic blue shift in the absorption spectrum ^1, 2^ and fluorescence quenching of up to 40-fold ^3^. The dimer to monomer transitions of multiple rhodamine dyes have measured dissociation constants of several hundred micromolar (supplementary table SI) and their properties have been described in terms of exciton theory ^2^.

Tetramethylrhodamine (TMR) is frequently used as a probe of biological processes ^2, 4–9^. Recently, TMR self-quenching has been extended to report on changes in proteins at single molecule resolution – to create a sensor for single adenosine 5’-diphosphate (ADP) molecules ^10^, to monitor conformational change of the activation loop in protein kinases ^11, 12^ and to monitor the dissociation of sliding clamps in DNA replication ^13^. TMR is expected to provide minimal disturbance to the physical behaviour of these relatively large biomolecules and – due to the expected shorter distance dependence of dimer-associated quenching compared with Förster resonance energy transfer (FRET) – has the potential to give detailed insight into very small distance changes or fast motions. TMR self-quenching has an additional advantage over FRET in that protein samples require labelling with a single dye at multiple sites, rather than a donor/acceptor dye pair. Nevertheless, the potential of TMR can only be fully realised if the distance-dependence and underlying timescale of the observed TMR signal change are known.

Previous studies of TMR have focussed on the affinity and orientation of dye dimerization ^5, 14, 15^. In this communication, we use dye-labelled model peptides to determine the distance dependence, timescale and process of TMR self-quenching. We determine that TMR is fully quenched when the average dye-dye distance is less than ~12 Å and that the observed rate constant for dye-dye interaction is fast. We expect that our results will enable us and others to use this phenomenon with confidence as a probe of increasingly subtle conformational change in biological molecules.

## Results and Discussion

### Developing a set of helical polyproline standards

In order to measure the distance-dependence of TMR self-quenching, we took a similar approach to Stryer and Haugland in their classical verification of Förster’s resonance energy transfer ^16^. We created a series of helical polyproline standards in which the terminal dye-labelled cysteine residues were separated by 0, 3, 5, or 6 proline residues (referred to throughout as 0P, 3P, 5P and 6P respectively; Figure 1A). Polyproline oligomers are often considered to be rod-like, although previous reports have noted the potential of cis-proline isomers to cause helical kinking ^17^ and of chain flexing in long polypeptides ^18, 19^. Neither of these caveats influences to our system: our samples fall within the range for which chain flexing can be excluded ^19^ and the probability of obtaining an all trans polyproline helix for the longest helix used in our work (*n* = 6) is between 81% and 90% (depending on how isomerisation of the terminal prolines is considered) ^20^.

**Figure 1:**
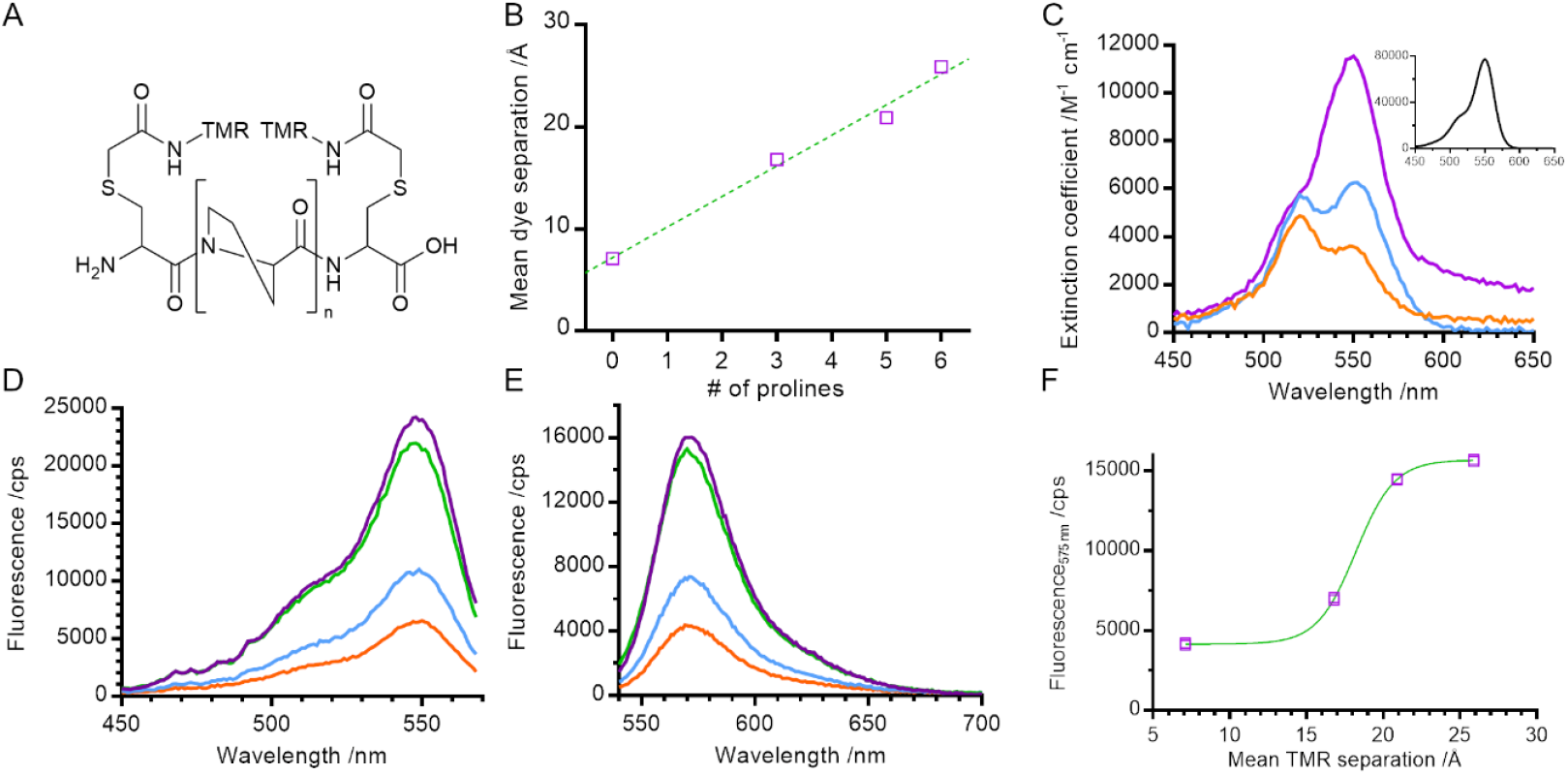
Physical and steady state spectral properties of polyproline helices (A) General structure of TMR labelled helices (n = 0, 3, 5 or 6 proline residues). (B) Mean dye separation, as calculated by FPS software. (C) UV-Vis absorbance of 0P, 3P and 6P. (inset) UV-Vis absorbance of hydrolysed TMRIA (ie free dye). (D) Steady state fluorescence excitation and (E) emission spectra of the polyproline length series. (F) Fluorescence emission at 575 nm against dye separation. Orange – 0P, blue – 3P, green – 5P, purple – 6P.

We built structural models of our helices and used FRET-restrained positioning and screening (FPS) software to determine the dye-accessible volume and mean dye positions ^21, 22^ (Figure 1B and supplementary figure S1). The distance between mean dye positions varied linearly with number of proline residues, with an approximate 3 Å increase in distance for each additional proline residue, consistent with the expected rise per residue for type II polyproline helices ^23^.

### Absorption spectra indicate dimer formation in short helices

For small electronic overlap, exciton theory states that the energy levels of an excited state dimer split in comparison with that of the excited state monomer (supplementary figure S2)^24^.

Experimentally, this results in a shift in the absorption spectrum of the molecule, the direction and magnitude of which is dependent on the orientation of the monomer units within the dimer ^24, 25^. This makes UV-Vis absorbance spectroscopy a probe of dimer geometry and of the monomer-dimer transition.

We measured the absorption spectra of our set of helical standards. The absorption spectrum for 6P had a peak at 550 nm and a shoulder at 520 nm, very similar to that of a TMR monomer (Figure 1C main panel & inset). Consistent with the exciton framework and an H-dimer in which the transition dipoles of the two monomer units are parallel ^2, 24, 25^, the main absorption peak for 0 P underwent a blue shift to 520 nm, overlapping with the vibronic shoulder of the TMR monomer and giving rise to an apparent reversal of intensities of the two peaks. This is similar to measurements of concentrated solutions of other rhodamines ^25, 26^, and is consistent with the NMR structure for a dimer of 6’ TMR-labelled peptide ^5^. The absorption spectrum for 3P is intermediate between the spectra of 6P and 0P, indicating a mixture of monomer and dimer species.

### Dimer formation quenches steady state fluorescence

We next characterised the steady state fluorescence of the polyproline length series. As the length of the helix decreased, the intensity of the fluorescence emission also decreased (Figure 1E). This suggests that the formation of a TMR dimer (evidenced by the absorption spectra) creates a complex that is less fluorescent than when the TMR molecules are held apart.

In order to determine the spectral properties of the species giving rise to fluorescence quenching we measured the fluorescence excitation spectra of our helices (Figure 1D). In contrast with a true absorption spectrum, a fluorescence excitation spectrum indicates the absorption spectrum of species which result in the emission of a photon at a specified wavelength. In our samples, the fluorescence excitation spectra of all helices resembled the absorption spectrum of a TMR monomer (main peak at 550 nm, vibronic shoulder at 520 nm). No blue shift – characteristic of dimer formation – was detected, even in samples where the absorbance measurements indicate dimer formation. This indicates that the TMR dimer species is either weakly fluorescent or non-emissive and that quenching of the net fluorescence signal reflects a reduction in the population of TMR monomer. This is consistent with the exciton model where rapid internal conversion between excited dimer singlet states prevents a radiative (fluorescence) emission from the permitted excited state back to ground state and instead promotes formation of the dye triplet state via intersystem crossing ^24^.

### TMR self-quenching monitors distance changes over the range 15 – 20 Å

In order to determine the distance-dependence of TMR fluorescence we plotted fluorescence emission at 575 nm against mean dye separation (*ie* distance; Figure 1F) and fitted the data to a sigmoid curve with a Hill coefficient of 1. Dye fluorescence increased sharply over the range 15 Å to 20 Å with a midpoint of 18 ± 1 Å (fitted value ± fitting error). This defines the distance range over which TMR self-quenching can be used to probe conformational change of biomolecules.

### Time-resolved fluorescence

In order to gain further information on the processes underlying the decrease in fluorescence emission intensity, we measured the fluorescence lifetime decay of our polyproline length series (Figure 2A). We used the instrument response function (IRF) to perform deconvolution fitting of our data and determined that each curve was best described by three exponential terms. Using a global fit (*ie* sharing the lifetime of each component across all helices), we determined the fitted decay lifetimes of our samples to be 115 ± 6 ps, 2.2 ± <0.01 ns and 16.3 ± 0.5 ns.

**Figure 2:**
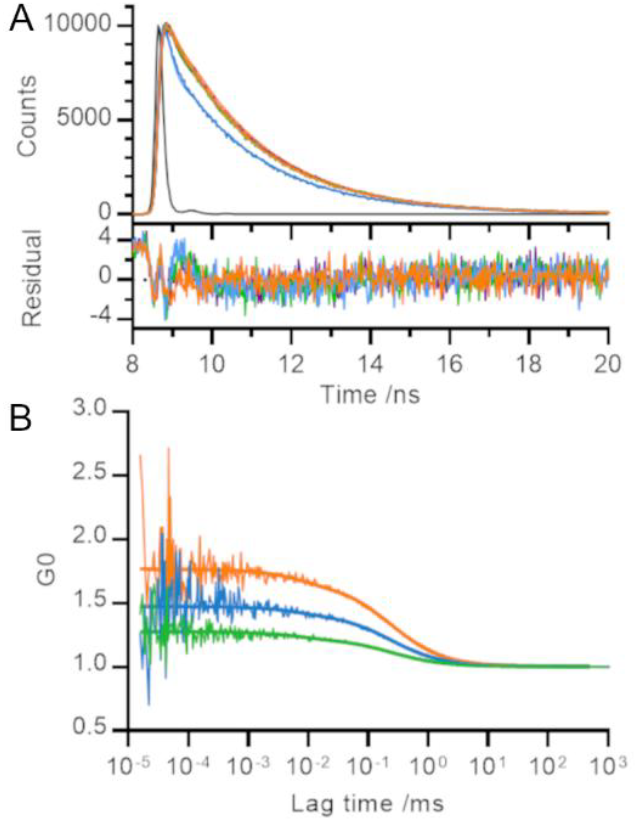
Temporal measurements of polyproline helices. (A) Time resolved fluorescence (time correlated single photon counting) and (B) Fluorescence correlation spectroscopy. Raw data and global fit (detailed in main text) shown. Orange – 0P, blue – 3P, green – 5P, black (A only) – instrument response function.

Despite the visible differences between the decay curve for 3P and those for rest of the length series, each decay trace was dominated by the fastest two phases (115 ps and 2 ns) and less than 0.5% of the trace arose from the slowest phase (16 ns; supplementary table SII). Our results are consistent with TMR lifetimes in protein environments measured elsewhere ^3, 5, 13, 15^. We converted the relative amplitudes of the three fitted phases into fractional populations (see supplementary Information for more details) to determine the underlying ratio of molecules in each environment to be approximately 10,000 : 1,000 : 1.

Unlike Donaphon and colleagues ^3^, we cannot assign the fast (115 ps) decay to quenching interactions between two dye molecules in close proximity since we observe this component in samples where the TMR dyes are unable to form a dimer. This is consistent with a description of the TMR-TMR interaction using exciton theory (in which dimerisation leads to formation of an excited state from which radiative decay is forbidden) rather than non-radiative return to ground state (*eg* by contact-induced Dexter quenching within the dimer; supplementary figure S2).

### Determining the timescale of TMR dimerization

TMR is an attractive dye to use a probe of protein dynamics. However, for this application to be implemented with confidence, it is important to characterise the timescale of formation / dissociation of the TMR dimer itself: no probe can be used to report accurately on the dynamics of a process occurring on a similar or faster timescale compared with the intrinsic fluctuations in the probe signal.

We set out to measure the timescale of TMR dimerization using our polyproline standards and fluorescence correlation spectroscopy (FCS). Previous FCS experiments involving TMR dimers have observed microsecond processes which were attributed to both protein dynamics ^27^ and potential photophysical properties of TMR ^28^. Our measured fluorescence correlation curves for 5P, 3P and 0P were similar to that for free TMR dye and showed no evidence of quenching dynamics within the timescales probed by the experiment (tens of ns up to the diffusion time of the helices through the confocal volume; Figure 2B). A global fit of these curves revealed a diffusion time of 0.24 ± 0.03 ms and a triplet lifetime of 7 ± 4 μs. The amplitude of the microsecond lifetime increased with laser power, confirming its assignment to the triplet state. We therefore conclude that the timescale of TMR dimerization must occur on a timescale faster than that probed by our FCS experiments (τ ≤ 100 ns), or one bounded by the helix diffusion time and the slowest process reported in the literature measured by TMR self-quenching (*ie* 0.24 ms ≤ τ ≤ 300 ms) ^10–12^.

The amplitude of fluorescence correlation curves contains information on the average number of fluorescent particles within the confocal volume. Smaller FCS amplitudes (lower G0) indicate an increase in the concentration of fluorescent particles. As the distance between TMR molecules decreased across the length series 5P → 3P → 0P, the number of apparent fluorescent particles found in solution decreased, even though the overall concentration of polyproline was held constant, consistent with our other measurements (Figure 2B).

### Comparison to FRET

The most common fluorescence method used to measure conformational dynamics of biomolecules is FRET, which is sensitive to changes on the nanoscale. Our results show that TMR is sensitive on a shorter length scale, and can thus report on dynamics or conformational change where conventional FRET pairs would be insensitive. TMR is also physically smaller than many FRET pairs, thus reducing the average distance between dye molecules on a labelled biomolecule and reducing the likelihood that the dye itself perturbs the behaviour of the biomolecule being measured.

One example where independent measurements of TMR self-quenching ^11, 12^ and conventional FRET pairs ^29^ have been used to monitor the same conformational change is on the activation loop of the protein kinase Aurora-A. We have compared these methodologies by calculating the expected distance between dye molecules for a TMR dye pair and for the FRET pair Alexa488/Alexa568 (supplementary table SIII and supplementary table SIV). We have then used the experimental results from single molecule TMR experiments ^11^ to predict the experimental bulk FRET distances ^29^ demonstrating excellent agreement between the two methodologies (supplementary table SV).

## Conclusion

In summary, we have used polyproline helices as spacers of defined length to determine the distance-dependence, timescale and process of TMR self-quenching. TMR self-quenching occurs over the distance range 15 – 20 Å, with maximal quenching occurring when dyes are closer than ~10 Å. Quenching occurs by a static process involving formation of a ground state dimer with a distinct absorption spectrum to that of monomer dye.

Our experiments here, and single molecule experiments on protein samples by us ^11, 12^ and others ^10^, indicate that two TMR molecules in close proximity are not completely dark and instead exhibit weak fluorescence. The emission of this weak fluorescence has similar spectral properties to unquenched TMR and a similar fluorescence lifetime. We hypothesise that this weak fluorescence reflects a time-average of dyes quickly interconverting between a dark dimer and two monomers close in space but incorrectly orientated for formal dimerization. This explanation is consistent with the results here, with exciton theory (in which splitting the energy levels within the excited state dimer depends on both the orientation and proximity of constituent monomers) and provides evidence that dye molecules can rotate relative to the protein to which they are tethered. Single molecule studies on protein samples indicate that photobleaching of either dye prevents quenching ^10–12^.

Formally, our measurements indicate that TMR dimerization either occurs faster than the timescale probed by our FCS experiments (τ ≤ 100 ns), or the region 1 – 300 ms. For consistency with our explanation of a weakly fluorescent species, we consider a fast process more probable.

Our measurements benchmark the use of TMR self-quenching as a probe of conformational change in biomolecules. In particular, they validate the use of self-quenching to probe conformational change in situations where protein size or geometry mean that dye labelling sites are constrained to be close in space, or where preparing a sample labelled with a donor/acceptor dye pair is unfeasible. Going forwards our measurements will enable researchers to make informed decisions on the choice of dye labelling sites under these (and other) conditions, and also to set bounds on the timescale of dynamic processes which can be monitored by this technique. Overall, our work provides the information necessary to enable TMR self-quenching to be used as a probe of protein conformational change complementary to traditional FRET measurements with confidence.

## Acknowledgements

CAD would like to thank Robert Kelsh for helpful discussions. This research was funded by the Wellcome Trust, AstraZeneca UK Ltd (RSRO_P71752 to CAD) through the Astra Zeneca/Imperial College London Innovation Fund, the National Institute for Health Research (NIHR) Biomedical Research Centre based at Imperial College Healthcare NHS Trust and Imperial College London (RSRO_P71816 to CAD) and BBSRC (JF20607/2 to LY). The views expressed are those of the author(s) and not necessarily those of the NHS, the NIHR or the Department of Health. A CC BY licence is applied to the Author Accepted Manuscript.

Experimental data for the figures in this paper, including models of the polyproline helices, are deposited in Zenodo (DOI: 10.5281/zenodo.12801405).

## Notes

https://doi.org/10.5281/zenodo.12801405

### Competing Interest Statement

The authors have declared no competing interest.

### Summary of Updates

Remeasurement of the absorption spectra in Figure 1C at lower sample concentrations. Minor amendments to the text for clarity. Supplemental information updated.

