## Supplementary information for "Tetramethylrhodamine self-quenching is a probe of conformational change on the scale of 15 – 25 Å"

### Abstract

Tetramethylrhodamine (TMR) is a fluorescent dye whose self-quenching has been used as a probe of multiple biological phenomena. We determine the distance-dependence of self-quenching and place bounds on the timescale of TMR dissociation. Our results validate fluorescence self-quenching as an alternative to FRET and enable future assays to be designed with confidence.

### Supplementary material

*Supplementary table S I: Reported dissociation constants for rhodamine dimers*

| Dye | Structure | $K_d$ /μM | Data source |
|---|---|---|---|
| Rhodamine 6G | 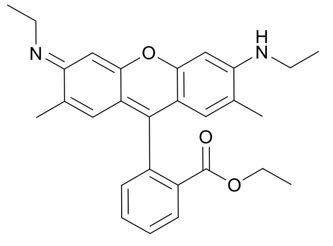 | 160 | [1] |
| | | 174 | [2] |
| | | 380 | [3] |
| | | 390 | [4] |
| | | 420 | [3] |
| | | 590 | [5] |
| | | 990 | [6] |
| Rhodamine B | 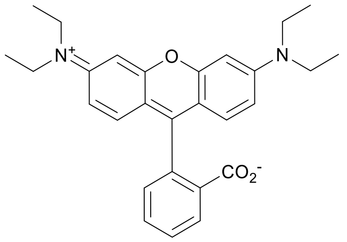 | 670 -770 | [3] |
| 5′ Tetramethyl rhodamine (5′ TMR) | 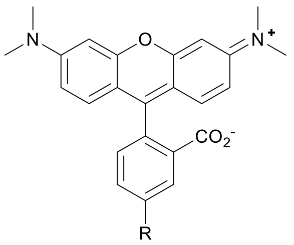 | 137 | [7] |
| 6′ Tetramethyl rhodamine (6′ TMR) | 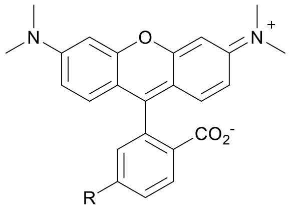 | 56 | [7] |

*Supplementary table S II: Relative percentage of each phase from global fitting of lifetime data*

| Sample | Rel % of 115 ± 3 ps | Rel % of 2 ± 5 ns | Rel % of 16 ± 5 ns | Avg. lifetime / ns |
|---|---|---|---|---|
| 6P | 26.7 [a] | 72.9 [b] | 0.4 | 1.69 [c] |
| 5P | 35.8 [a] | 63.8 [b] | 0.4 | 1.49 [c] |
| 3P | 58.2 | 41.5 | 0.2 | 1.01 |
| 0P | 34.6 [a] | 65.1 [b] | 0.3 | 1.52 [c] |

[a]The average contribution of the fast amplitude for these three helices is 32 ± 5 %
[b]The average contribution of the 2 ns amplitude for these three helices is 67 ± 5 %.
[c]The overall average lifetime for these three helices is 1.6 ± 0.1 ns.

*Supplementary table S III: Measured distances for TMR-labelled Aurora-A kinase*

| | Distance /Å<br>Active conformation bound to ADP (1OL7) | Active conformation bound to ADP and protein activator TPX2 (1OL5) | Inactive conformation bound to MLN8054 (2WTV) |
|---|---|---|---|
| $C_\alpha$-$C_\alpha$ [a] | 36.5 | 42.0 | 15.1 |
| $R_{mp}$ [b] | 43.9 | 50.4 | 18.2 |
| $\langle R_{dye\text{-}dye} \rangle$ [c] | 44.3 | 50.9 | 19.2 |

[a] Residue 224 to 283 (labelling positions used in [8, 9])
[b] Distance between mean positions of dyes (calculated using FRET-restrained positioning and screening (FPS) software [10, 11])
[c] Mean distances between dyes (calculated using FPS software [10, 11])

*Supplementary table S IV: Expected FRET efficiencies for Alexa488/Alexa568 dye pair*

| | Active conformation bound to ADP (1OL7) | | Active conformation bound to ADP and protein activator TPX2 (1OL5) | | Inactive conformation bound to MLN8054 (2WTV) | |
|---|---|---|---|---|---|---|
| | K224/S283 labelling sites [8, 9] | S284/L225 labelling sites [12] | K224/S283 labelling sites [8, 9] | S284/L225 labelling sites [12] | K224/S283 labelling sites [8, 9] | S284/L225 labelling sites [12] |
| $C_\alpha$-$C_\alpha$ /Å | 36.5 | 36.8 | 42.0 | 41.0 | 15.1 | 14.6 |
| $R_{mp}$ /Å [a] | 52.8 | 54.1 | 56.3 | 56.3 | 21.0 | 20.9 |
| $\langle R_{DA} \rangle$ /Å [b] | 55.5 | 57.1 | 59.4 | 59.3 | 27.0 | 26.6 |
| $\sigma_{DA}$ [c] | 7.1 | 7.1 | 7.9 | 7.2 | 10.8 | 10.1 |
| *E* /% [d] | 65 | 61 | 56 | 57 | 97 | 98 |
| $\langle R_{DA} \rangle_E$ /Å [e] | 55.9 | 57.4 | 59.5 | 59.3 | 34.0 | 32.9 |

[a] Distance between the mean positions of donor and acceptor dyes within the accessible volume
[b] Mean distances between donor and acceptor dyes within the calculated dye-accessible volume
[c] Fitted width of the $\langle R_{DA} \rangle$ distribution
[d] Expected FRET efficiency (based on calculated mean positions and reported dye $R_0$)
[e] FRET-averaged apparent distance between donor and acceptor molecules:

$$\langle R_{DA} \rangle_E = R_0 \left( \frac{1}{\langle E \rangle} - 1 \right)^{\frac{1}{6}}$$

All parameters detailed in footnotes calculated using FPS software. More details can be found in software notes and publication [10, 11].

*Supplementary table S V: Comparison of TMR and FRET measurements of conformational change in Aurora-A kinase*

| Experimental condition [a] | Measured population of active conformation /% [b] | Calculated distance /Å [c] | Experimental distance / Å [d] |
|---|---|---|---|
| Apo kinase | 77 | | 51.5 |
| ADP | 78 [e] | 52 | 54.7 |
| TPX2 | 86 | 56 | 56.5 |
| TPX2 / ADP | 88 [f] | 56 | 59.5 |

[a] All measured using phosphorylated kinase domain

[b] From Table I, reference [9]

[c] $r_{calc} = popn_{active} \cdot \langle R_{DA} \rangle_{E\_active} + (1 - popn_{active}) \cdot \langle R_{DA} \rangle_{E\_inactive}$ where $r_{calc}$ is the calculated value for the experimentally observed FRET distance, $popn_{active}$ is the population of molecules in the active conformation (expressed as a fraction), $\langle R_{DA} \rangle_{E\_active}$ is the FRET-averaged apparent distance between donor and acceptor molecules for the active conformation (Supplementary table S IV, column 3 or 5) and $\langle R_{DA} \rangle_{E\_inactive}$ is the FRET-averaged apparent distance between donor and acceptor molecules for the inactive conformation (Supplementary table S IV, column 7).

[d] From Figure 2c, reference [12]

[e] ADP not used as a ligand for single molecule measurements, so value in presence of saturating quantities of AMP-PNP used instead

[f] Value is for kinase in presence of saturating quantities of TPX2, AMP-PNP and substrate peptide kemptide

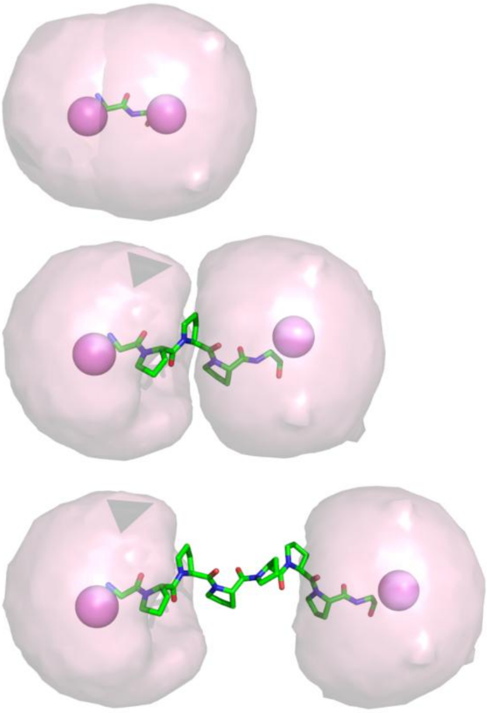

*Supplementary figure S 1: Dye-accessible volumes for polyproline helices with zero, three and six proline residues. The dye clouds overlap for 0P, are close to overlap for 3P and have no overlap for 6P.*

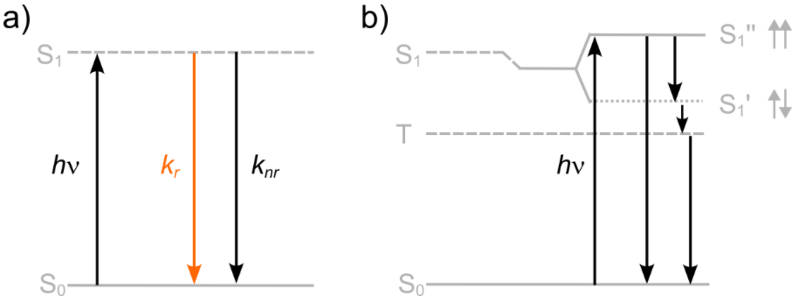


*Supplementary figure S 2: Energy diagrams for TMR monomer and exciton dimer. a) TMR monomer. In the monomer, absorption of a photon leads to excitation of an electron from ground state ($S_0$) to an excited state ($S_1$). The electron returns to ground state either through emission of light (fluorescence, radiative transition; $k_r$) or by non-radiative processes such as thermal vibration ($k_{nr}$). b) TMR dimer. In the TMR exciton dimer, the energy level of the excited state has split in two [13]. The in-phase dipole interaction raises the energy of the excited state ($S_1''$) compared to the monomeric energy level for $S_1$, the out-of-phase dipole arrangement lowers the energy of the excited state ($S_1'$). Arrows to the right of each energy level indicate dipole phase relation. Transitions from ground state to exciton state $S_1'$ are forbidden, while transitions to $S_1''$ are permitted. The increase in the energy difference for excitation in the dimer ($S_1''$-$S_0$) compared with monomer ($S_1$-$S_0$) gives rise to the blue shift observed in the absorption spectrum upon dimerisation, and confirms that the two transition dipoles are parallel in the dimer (in-line transition dipoles lead to different splitting and a red shift in absorption). Upon absorption of a photon, rapid internal conversion between singlet states prevents emission as fluorescence ($S_1'$ – $S_0$ transition is forbidden), leading to dissipation of energy either through thermal vibrations or through intersystem crossing and return via the triplet state. Panel b) redrawn from reference [13].*

### Experimental details

#### Modelling inter-dye distance

The PolyprOnline database https://www.dsimb.inserm.fr/dsimb_tools/polyproline/[14] was used to identify PDB ID 1OWL (apo DNA photolyase from *Anacystis nidulans* [15]) as a high (1.8 Å) resolution x-ray structure containing an extended region of polyproline helix (PKPTPVATP – residues 160-169 – identified as a polyproline helix by all algorithms). The 3D coordinates of this sequence (and the two flanking residues) were extracted from the record, and the central residues mutated to consecutive proline residues using the mutagenesis wizard within PyMOL [16]. All new residues fitted easily into the helix and no geometric clashes were observed. Helices were truncated to match the length of experimental samples (0, 3, 5 and 6 proline residues) and flanking N- and C- terminal residues mutated to glycine (not cysteine). This ensured that flexibility and rotation in the $C_\alpha$–$C_\beta$ and $C_\beta$–S bonds were included in the subsequent modelling.

FRET-restrained positioning and screening (FPS) software [10, 11] was used to model the dye-accessible volume for each construct and to determine the mean dye position. This software takes a (static) pdb structure, information about the geometry of the dye and the way that the dye is attached to the biomolecule, and calculates the volume of space that the dye can occupy assuming an equal probability at each position. Using this accessible volume, it reports the mean dye position. FPS software does not consider molecular interactions such as dye stacking.

Dyes were modelled as an ellipsoid of 7.1 × 4.3 × 1.8 Å which were the width, height and thickness respectively of the planar xanthene ring system estimated using an energy-minimised model within Chem3D (PerkinElmer). The linker length was measured from the $C_\alpha$ carbon of the terminal glycine to the xanthene moiety of the dye. The length of the linker was determined to be 8.3 Å with a width of 4.5 Å. Overlap of the dye clouds was observed for 0P sample only. Mean dye separation was measured in PyMOL as the distance between mean dye positions.

#### Experimental samples

Polyproline helices (1 mg scale, >98% purity and capped at both N- and C- termini) were purchased from Peptide Synthetics (Peptide Protein Research Ltd, Fareham, UK). Peptide sequences were Ac-C(TMRIA)C(TMRIA)-NH2 (referred to in main text as 0P), Ac-C(TMRIA)PPPC(TMRIA)-NH2 (referred to as 3P), Ac-C(TMRIA)PPPPPC(TMRIA)-NH2 (referred to as 5P), and Ac-C(TMRIA)PPPPPPC(TMRIA)-NH2 (referred to as 6P). C(TMRIA) is a cysteine residue reacted with 5-tetramethylrhodamine iodoacetamide to result in covalent attachment of the dye to the peptide through the thiol.

Peptide lengths were chosen so that our measurements would sample the expected distance range of TMR self-quenching (~15-21 Å) [17].

Polyproline helices have previously been used to verify the $1/r^6$ dependence of FRET [18], distance-dependent FRET at the single molecule level [19] and, very recently, super-resolution measurements of short polyproline helices have shown that helices shorter than 15 residues to give rise to a single distribution of end-to-end distances [20]. Polyproline helices have been shown to be extremely robust to denaturation by detergent, heat, pH and chaotropic agents [21]. Together, these data give us confidence that the structure of our (very short) dye-labelled helices remains unchanged by dye labelling.

#### Absorbance and steady state fluorescence spectroscopy

Absorption measurements were made using a Cary 60 spectrophotometer (Agilent Technologies) using a quartz absorption cuvette (Hellma Analytics) with a pathlength of 10 mm. Measurements were made at room temperature using a concentration of 1 μM peptide in 50 mM Tris pH7.5, 200 mM NaCl, 5 mM $MgCl_2$, 10% glycerol. The absorption spectrum of buffer alone was subtracted from all spectra.

We reiterate that the sample concentration above (1 μM) refers to the concentration of double-labelled peptide. When determining the extinction coefficient of TMR, absorption values were normalised using the concentration of *dye* (ie 2 μM – double that of peptide – due to the presence of two dye molecules per helix). We note that the extinction coefficient ($\varepsilon_{550}$) for 6P ($\varepsilon_{550,\,6P}$ = 12,000 $M^{-1}$ $cm^{-1}$) is much smaller than our measurement of the extinction coefficient for free hydrolysed dye ($\varepsilon_{550,\,free\,dye}$ = 77,000 $M^{-1}$ $cm^{-1}$), the latter being consistent with that reported elsewhere [22].

Steady state fluorescence measurements were made with a FluoroMax-4 (HORIBA) spectrofluorometer and quartz micro fluorescence cuvettes (Hellma Analytics) with a light path of 10 mm × 2 mm. Emission spectra were measured using an excitation wavelength of 532 nm with an excitation slit width of 5 nm and emission slit width of 2 nm. Excitation spectra were collected using a detection wavelength of 575 nm with an excitation slit width of 2 nm and emission slit width of 5 nm. Measurements were made at room temperature in 10 mM phosphate pH 7.0. The sample concentration for fluorescence measurements was 20 nM. Under these conditions we expect inner filter effects and inter-molecular self-quenching to be negligible.

**Time resolved fluorescence spectroscopy**

Time resolved fluorescence was obtained using a DeltaFlex (Horiba) Time-Correlated Single Photon Counting system using micro fluorometer cells (Starna scientific) constructed of special optical glass with a path length of 5mm. Excitation was provided by a 467 nm NanoLED (Horiba) with a repetition rate of 1MHz. While this is not a typical excitation wavelength for TMR, the NanoLED emission was broad enough that it could still provide inefficient excitation of TMR. For this reason, the concentration of polyproline samples used for time resolved measurement (10 μM) was higher than in our steady state measurements (20 nM). Even with the increased concentration the count rate was still low and so the acquisition time was increased in order to achieve a peak present of 10,000 counts and therefore adequate signal to noise.

Fluorescence was filtered using a 550 nm long pass filter before being detected at 575 ± 4 nm with a bandpass filter. The count rate was kept ≤ 1%. All measurements were made at 23 °C in 10 mM phosphate buffer pH 7.0 with a sample concentration of 10 μM. Data was analysed using DAS-6 (Horiba) software.

Fluorescence lifetime data was fit using DAS-6 (Horiba) software. Data was globally fit to three lifetime components, sharing the lifetime across all traces. Data fitting with fewer or more lifetime components did not provide better fitting of the data based on the chi squared value and quality of residuals.

The relative amplitudes reported are the normalized pre-exponential factors calculated by

$$f_i = \frac{A_i}{\sum A_i} \tag{S1}$$

where $f_i$ is the fractional (relative) amplitude for each phase and $A_i$ is the fitted pre-exponential factor (amplitude).

**Fluorescence correlation spectroscopy**

Fluorescence correlation spectroscopy (FCS) measurements were performed on a home-built confocal setup. Excitation was provided by a tuneable argon ion laser (35LAP321-230, Melles Griot,

USA) set at 514 nm. In this system, the beam is passed into an inverted microscope (Nikon Eclipse TE2000-U) before being focused 6 µm into the sample with a high numerical aperture (60× 1.45 NA oil) objective. The laser intensity at the point of entry into the microscope for the data presented in Figure 2 of the main paper was 210 µW. These measurements were repeated at 600 µW to confirm of assignment of the microsecond lifetime to the triplet state.

Fluorescence emission from the sample is passed through a pinhole before being split using a 50/50 beam splitter. Both beams pass through identical bandpass filters (575-50m, Chroma), before being detected using identical avalanche photodiode detectors (SPCM-AQR-14). The two detectors are connected to a digital hardware correlator (flex02-01d/c). Signals from each detector are cross correlated, which produces a pseudo autocorrelation curve thus avoiding artifacts due to the dead time of the detectors.

FCS measurements were performed using 5nM TMR-labelled polyproline helices in 50 mM Tris-HCl pH 7.5, 200 mM NaCl, 5mM $MgCl_2$, 10% glycerol, 1% DMSO, 0.3 mg $ml^{-1}$ BSA. Under these conditions we expect inner filter effects and inter-molecular self-quenching to be negligible.

Fitting of the FCS traces was performed in Origin (OriginLab) using equation (S2), which accounts for the (2D) diffusion of the species through the confocal volume, with diffusion time ($\tau_D$), and a fraction (F) of the molecules being in the triplet state with a corresponding triplet lifetime ($\tau_F$).

$$G(\tau) = \frac{G(0)}{1+\frac{\tau}{\tau_D}}\left(1-F+F\,e^{-\frac{\tau}{\tau_F}}\right)+C \qquad \text{(S2)}$$

### Converting fluorescence lifetime amplitudes into proportion of molecules

Our fluorescence lifetime measurements were best fit by a decay curve with three lifetimes:

$$decay = f_1\tau_1 + f_2\tau_2 + f_3\tau_3 \tag{S3}$$

where $f_i$ is the fractional amplitude for phase *i*, $\tau_i$ is the lifetime for phase *i*, and *decay* is the observed total fluorescence lifetime decay. $\sum_i f_i = 1$. This indicates that our TMR dye molecules experience three distinct environments, each giving rise to a different decay lifetime.

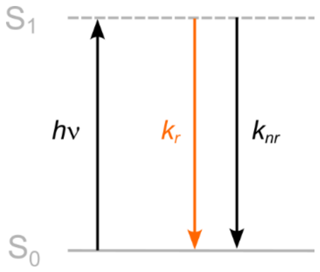


*Figure 1: Energy diagram for simple fluorescence. Absorption of a photon leads to an electron being excited from ground state ($S_0$) to an excited state ($S_1$). The electron can return to ground state via fluorescence emission (orange; with rate constant $k_r$) or via non-radiative processes such as thermal vibrations (with combined rate constant $k_{nr}$).*

Each decay lifetime, $\tau_i$, is the reciprocal of the associated rate constant for return from the excited state ($S_1$) to ground state ($S_0$) in that environment, $k_i$. $k_i$ is the sum of the rate constants for radiative ($k_r$) and non-radiative ($k_{nr}$) relaxations from the excited state (Figure 1). Written mathematically,

$$\tau_i = \frac{1}{k_i} = \frac{1}{k_r + k_{nr_i}} \tag{S4}$$

Since the electronic transition is between the same excited state and ground state ($S_1$ to $S_0$) in all environments, the rate constant for radiative decay, $k_r$, is constant across all environments. Different fluorescence lifetimes are instead due to the presence of different non-radiative pathways in each environment (*i.e.* to different values of $k_{nr}$). This gives rise to the subscript *i* on $k_{nri}$ in equation (S4).

For any fluorescence emission, the probability that an individual transition gives rise to fluorescence is given by the probability that an electron returns to ground state via a radiative process compared with a non-radiative one. *i.e.*,

$$P(fluorescence) = \frac{k_r}{k_r + k_{nr}} \tag{S5}$$

As the net speed of non-radiative pathways increases – as the number of rapid non-radiative pathways increases – the value of $k_{nr}$ increases. As $k_{nr}$ becomes large compared with $k_r$, not only does the probability of a fluorescence during a transition decrease (S5), the contribution of $k_r$ to the overall rate constant $k_i$ also decreases:

$$k_r + k_{nr} \approx k_{nr} \text{ so } k_i \approx k_{nr} \tag{S6}$$

The total number of molecules in the sample, *N*, is made up of the number of molecules in each of the three environments indicated in equation (S3):

$$N = n_1 + n_2 + n_3 \tag{S7}$$

where $n_i$ is the number of molecules in environment *i*. The fraction of molecules in environment *i* is therefore given by $\frac{n_i}{N}$. (S8)

The fractional amplitude for the phase of a fluorescence lifetime decay depends on both the number of molecules in the environment giving rise to that decay and the probability that each electronic return to ground state gives rise to fluorescence. Putting equations (S8) and (S5) together, we can say that for each fractional amplitude, $f_i$, in equation (S3),

$$f_i = \frac{n_i}{N} \cdot \frac{k_r}{k_r + k_{nr_i}} \tag{S9}$$

Rearranging and substituting from (S4)

$$\frac{n_i}{N} = \frac{f_i}{k_r \tau_i} \tag{S10}$$

$$n_i \propto \frac{f_i}{\tau_i} \tag{S11}$$

$k_r$ and *N* are constants within any set of experimental traces, meaning that the ratio of the populations of molecules in each environment is exactly equal to the ratio of the experimental fractional amplitude divided by the lifetime:

$$n_1 : n_2 : n_3 = \frac{f_1}{\tau_1} : \frac{f_2}{\tau_2} : \frac{f_3}{\tau_3} \tag{S12}$$
